# Heterogeneous Effects Across Clonal Hematopoiesis of Indeterminate Potential (CHIP) Driver Mutations Challenges a Unified Model of CHIP

**DOI:** 10.64898/2026.11.24.768778

**Authors:** Matthew H Liu, Xinyu Sun, Lissette Gomez, Tianjie Gu, Makaela Mews, Yousef Mustafa, Anthony Griswold, William S Bush

## Abstract

Clonal Hematopoiesis of Indeterminate Potential (CHIP) is an age-associated expansion of hematopoietic stem cells harboring somatic mutations in the absence of overt malignancy. CHIP driver genes impart divergent effects on many biological processes, yet the mechanisms underlying these gene-specific effects, particularly in non-hematologic disease contexts, remain poorly understood. We analyzed 466 samples from the Multi-Ancestry Genomics, Epigenomics, and Transcriptomics of Alzheimer’s Disease dataset, and identified function-specific differences between CHIP categorizations. Uniquely, *DNMT3A* CHIP was associated with global hypomethylation, *STAG2* CHIP increased mutation burden, HSC differentiation CHIP strongly triggered adaptive immune response, and proliferative signaling CHIP induced coagulation and cell adhesion. These divergent functions differentially modulate inflammatory pathways and suggest investigating CHIP’s downstream effects through a driver gene function-based approach rather than treating CHIP as a single entity. Our multi-omics analyses provide such a framework to improve mechanistic interpretation of conflicting relationships between CHIP and diseases such as Alzheimer’s disease, cardiovascular disease, and cancer.

## Introduction

Clonal Hematopoiesis (CH) is an expansion of hematopoietic stem cell (HSC) clones harboring somatic mutations in the absence of malignant phenotypes with direct links to multiple age-related diseases, including hematologic cancers and cardiovascular disease.^1–12^ CHIP is defined as the presence of clonal hematopoiesis with somatic mutations occurring within individual hematological malignancy-associated (driver) genes at an allele fraction (VAF) ≥ 0.02.^13,14^ CHIP additionally confers pro-inflammatory effects that likely impacts progression of many non-hematological diseases.^15–19^ CHIP driver genes impart differing effects on multiple biological processes, including epigenetic regulation, genome stabilization, RNA splicing, HSC differentiation, and metabolic and proliferative signaling.^20–23^ Yet, the study of the downstream effects of CHIP mutations in these driver genes on an individual basis remains limited, and he heterogeneous mechanisms underlying CHIP may be a source of variability of observational reports in the literature. For example, CHIP is typically linked to worse outcomes in solid tumor patients,^24–28^ including thyroid and ovarian cancers, but *TET2* CHIP has been linked to improved immunotherapy in solid tumors.^29^ Liu et al. and Choi et al. have suggested CHIP could contribute to multiple neurodegenerative diseases,^30^ namely vascular dementia and amyotrophic lateral sclerosis, but also Alzheimer’s disease (AD).^31^ Yet, others have shown CHIP to be protective against Alzheimer’s,^32^ while other reports showed no correlation.^33^ Furthermore, CHIP has been shown to be generally contributory to acute kidney injury and CKD,^34,35^ but in some cohorts, there appears to be no relationship.^36^

To investigate the potential divergent effects of different forms of CHIP, we analyzed 466 samples from the Multi-Ancestry Genomics, Epigenomics, and Transcriptomics of Alzheimer’s Disease (MAGENTA) dataset,^37^ which contains high participant diversity (Supplementary Table 1) to supplement the limited CHIP research of non-White participants.^38–40^ We stratified participants by the likely functional impact of CHIP driver mutations, and observed how these endotypes differentially impact genomic stability, DNA methylation, transcriptional stability, cellular proliferation, immune activation, and systemic effects on the immune repertoire. We also observed how CHIP’s effects scale with clone size and genetic ancestry alters mutation frequency of individual CHIP driver genes.

Here, we propose a system for stratifying CHIP driver mutations based on driver gene function. We show these alterations to CHIP driver genes convey unique and specific effects that are often opposing, and therefore recommend that CHIP should be treated and studied as a collection of separate endotypes with divergent effects, rather than a single mechanism. Our results caution against making broad associations between CHIP and other diseases or phenotypes and generalizing the effects of one endotype to CHIP as a whole, due to clearly different and often opposing effects of different endotypes.

## Methods

### MAGENTA dataset

We analyzed 466 samples from the Multi-Ancestry Genomics, Epigenomics, and Transcriptomics of Alzheimer’s Disease (MAGENTA) dataset that had whole-genome sequencing (WGS) and complete data,^37^ containing participants aged 53-98 from African American, Puerto Rican, Peruvian, Cuban, and Non-Hispanic White ancestry. For each participant, we analyzed DNA Methylation assays, and bulk RNA-sequencing.

### CHIP Calling

Somatic mutations were identified from WGS data for each sample using the Genome Analysis Toolkit (GATK)’s *Mutect2* somatic variant caller.^41^ The reference genome used was the Homo_sapiens_assembly38.fasta along with the af-only-gnomad.hg38 germline resource and the 1000g_pon.hg38 panel of normals. To identify CHIP driver mutations, *Mutect2* was run on the regions surrounding canonical CHIP driver genes. Mutations were then annotated by gene using *ANNOVAR*.^42^ Quality control was performed using *BCFtools* to remove low fidelity variants. Mutations were then identified as “CHIP driver” mutations if they passed the putative CHIP whitelist filter.^43^ For genome-wide and CHIP region mutation burden, *BCFtools* was used to determine mutation count for each sample from WGS data. Significance between CHIP endotypes for mutation burden results were calculated using Welch’s t-test, with non-CHIP samples as the reference group.

### Methylation Analysis

Our MAGENTA study contains methylation levels for 854,672 CpG loci, with 708,062 CpG sites used for CHIP analysis. Methylation matrices were preprocessed using the *SeSAMe* R package with background correction and functional normalization.^44^ Cell type proportions (including CD4⁺ T cells, CD8⁺ T cells, B cells, natural killer cells, monocytes, neutrophils, and eosinophils) were estimated with the Houseman deconvolution method through the *EpiDISH* R package.^45^ Differentially methylated positions (DMPs) were identified using linear models implemented in limma, with empirical Bayes moderation to improve variance estimation across CpG sites. Models were adjusted for age, sex, ancestry, estimated cell type proportions, and batch effects. Statistical significance for DMPs was controlled for multiple testing using the Benjamini–Hochberg false discovery rate (FDR), with FDR < 0.05 considered significant. Kolmogorov-Smirnov (KS) tests were performed to determine the degree to which each CHIP endotype deviated from a reference probability distribution, and results were plotted in quantile-quantile plots. Differential global methylation significance was determined using the *globaltest* R package. CpG locations were annotated to genes using the *IlluminaHumanMethylationEPICanno.ilm10b4.hg19* R package. Methylation analysis results were displayed in volcano plots built using the *GGPlot2* R package. To assess biological relevance, significant DMPs were subjected to Gene Ontology (GO) enrichment analyses.^46–48^ Enrichment significance was evaluated using permutation-based testing with correction for multiple comparisons. Horvath age was calculated using the 353 CpG sites that form the Horvath epigenetic age clock (*methylclock* R package).^49^ Significance between CHIP endotypes for all methylation analysis results were calculated using Welch’s t-test, with non-CHIP samples as the reference group.

### Bulk RNA Analysis

60,668 protein-coding and non-coding genes were identified and used for differential RNA expression analysis. Statistical significance was determined again using Benjamini-Hochberg FDR correction, with FDR < 0.05 considered significant. Effect sizes were reported as log2 fold changes with corresponding confidence intervals. RNA expression analysis results were displayed in volcano plots built using the *GGPlot2* R package, or in heatmaps built using the *PHeatmap* R package. To assess biological coherence and mechanistic relevance, differentially expressed genes were subjected to GO biological pathway and gene-set enrichment analyses. Genes used for immune and cell proliferation programs were curated from RefSeq Gene Summary.^50^ For comparison between CHIP endotypes, Z-scores for each program were calculated for each endotype. Overlap analyses were conducted between genes linked to significant DMPs and genes identified as differentially expressed. From overlap analyses, a graph was constructed with endotype-program combinations with Z-scores greater than 2. Significance between CHIP endotypes for RNA expression analyses was calculated using Welch’s t-test, with non-CHIP samples as the reference group.

### RNA Splice Counting

RNA splice fractions were taken from bulk RNA sequencing as a fraction of total spliced reads divided by total reads for each sample.^51^ Reads were determined to be spliced using Samtools. Significance in splice count differences between CHIP endotypes was calculated using Welch’s t-test, with non-CHIP samples as the reference group.

## Results

### Classifying CHIP Endotypes Based on Known Mutant Gene Function and Metadata

To investigate the heterogeneity of CHIP in our dataset, we stratified CHIP by driver gene, clone size and ancestry. Participants with two or more CHIP mutations in the same driver gene were counted once, but participants with mutations in two or more CHIP genes were counted for all genes (Supplementary Table 2). We grouped CHIP driver genes based on function defined by their RefSeq Gene Summary, allowing for analysis of more rarely mutated CHIP driver genes. We observed that CHIP driver genes could be categorized into having effects on genome stability, epigenetic modulation, RNA splicing, hematopoietic cell (HSC) differentiation, and proliferative signaling (Figure 1A). We also grouped these major groups into whether they affected regulatory machinery (genome stabilization, and epigenetic regulation) or phenotypic adaptation (RNA splicing mediation, HSC differentiation and metabolic and proliferative signaling). The epigenetic regulators can broadly be subcategorized based on canonical function into genes that are either generally associated with increased gene expression (*TET2, SETD2, CREBBP, KDM6A*) or decreased gene expression (*DNMT3A, SETDB1, BCORL1*). Furthermore, the CHIP driver genes affect expression through either DNA methylation (*DNMT3A* and *TET2*) or histone modifications, while the ASXL family (*ASXL1* and *ASXL2*) and *SETDB1* were investigated independently of the histone modification CHIP genes due to their broad, genome-wide effects on chromatin remodeling,^52^ and larger sample size in our cohort, respectively. With CHIP genes involved in genome stabilization, *STAG2* and the Cohesin complex (*STAG2*, *PDS5B*, and *RAD21*) were investigated both together and individually due to large sample size and shared effects.^53^ *SF3A1*, *SF3B1*, *SRSF2*, *ZRSR2*, and *PRPF40B* are all involved in RNA splicing, and were further stratified into spliceosome components (*SF3B1* and *ZRSR2*) and splicing regulators (*SF3A1*, *SRSF2*, and *PRPF40B*). For CHIP genes that affect specific functions and pathways, *CSF3R*, *IKZF2*, *IKZF3*, and *RUNX1* all contribute to HSC differentiation into distinct hematopoietic cell types, and were thus grouped together. Finally, multiple genes affect cell proliferation, and again could do so by either increasing or decreasing proliferation. Genes that were part of the PI3K/AKT (*PTEN* and *PHIP*) or Ras/MAPK (*GNAS*, *NF1*, *GNB1*, and *CBL*) pathway were stratified for separate analyses as well. The PI3K/AKT pathway is the body’s primary metabolic cascade, while the Ras/MAPK pathway is the body’s primary mitogenic and proliferative cascade.^54^ Participant samples were stratified into large (VAF>0.1) and small (VAF<0.1) clones, as defined in previous literature,^55–57^ to investigate if larger clone size would exacerbate CHIP driver gene effects.

**Figure 1:**
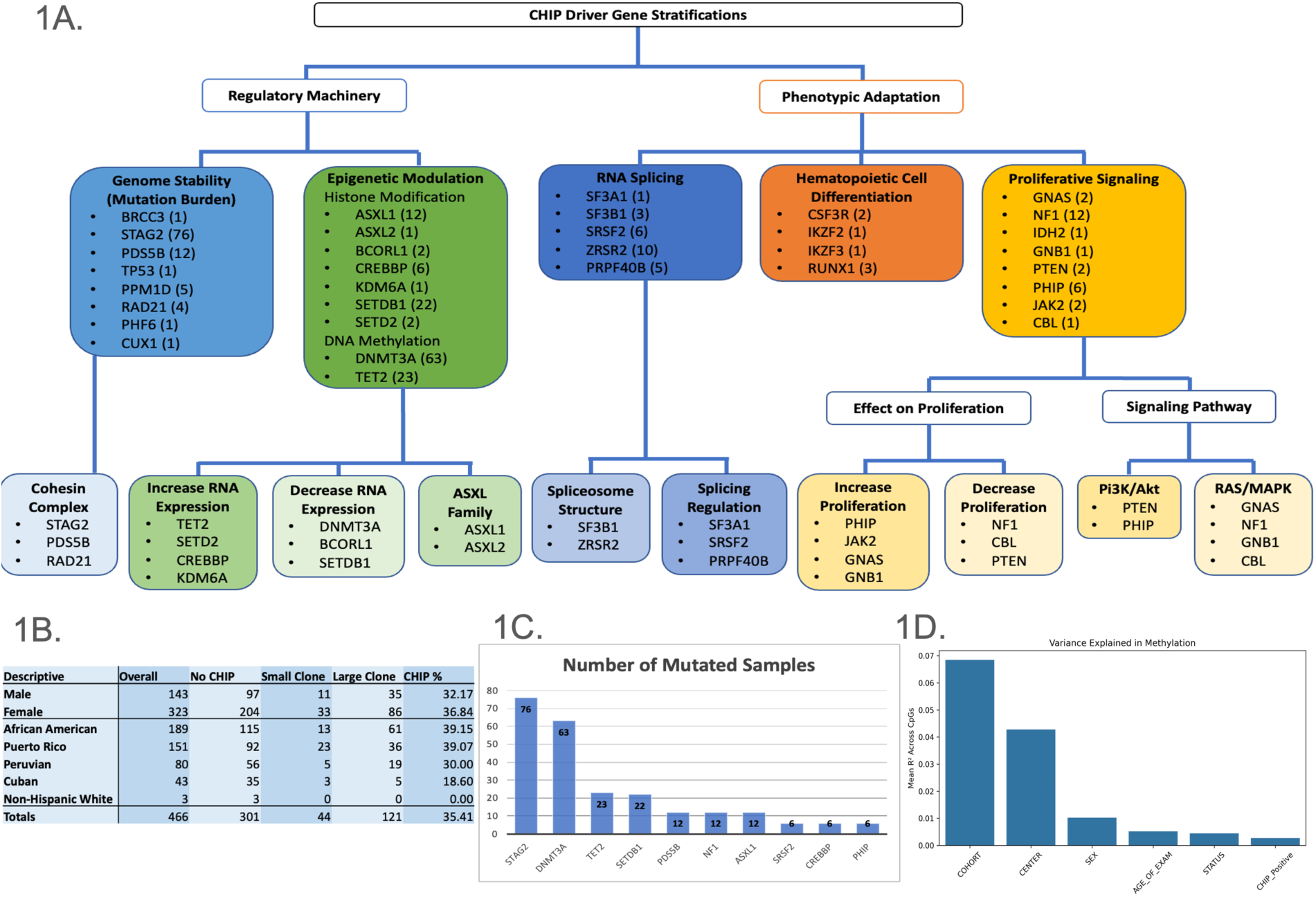
Stratifying CHIP by Driver Gene, Clone Size, and Ancestry. **a**, CHIP driver genes categorized by global or specific pathway effects, then by driver gene function, and further by either pathway or direction. The number of samples with each driver gene mutation is in parentheses. Samples with multiple mutation CHIP genes were counted for each gene, but samples with multiple mutations in the same CHIP gene were not double counted. **b**, Prevalence of non-CHIP, and large and small clone CHIP in Males vs. Females, and by ancestry. **c**, The most common CHIP driver mutations in the MAGENTA dataset. **d**, Coefficient of determination demonstrating variance in methylation explained by ancestry, center, sex, age, AD status and CHIP status.

Primary outcomes observed include total and CHIP region mutation burden, global methylation changes, RNA splice fraction, global RNA expression changes, cell type proportions, significantly differentially expressed genes, and differential immune program expression. For all outcomes, results for each CHIP endotype were calculated, but only significant results and results for endotypes whose canonical functions are directly related to the outcome were reported in the results section. For significantly differentially expressed genes, results for genes that are related to the same pathway as the CHIP gene or are part of an enriched GO pathway were reported.

The MAGENTA dataset consisted of 189 African American, 151 Puerto Rican, 80 Peruvian, 43 Cuban, and 3 Non-Hispanic White participants (Table 1B), all of whom have accessible whole-genome sequencing, bulk-RNAseq, and DNA methylation data. Of the 466 total samples, 165 (35.4%) had a mutation in at least one CHIP driver gene that passed the whitelist filter.^43^ Of those 165 samples, 121 were large clones and 44 were small clones. The most common CHIP driver mutations in our MAGENTA dataset were *STAG2* (76 samples), *DNMT3A* (60), *TET2* (23), and *SETDB1* (22) (Figure 1C). This somewhat aligns with the typical distribution of CHIP driver genes, however, *STAG2* is notably the most abundantly mutated gene in our dataset, which is generally less abundant in previously reported, predominantly White datasets.^58^ Additionally, when performing DNA methylation analysis, samples clustered by ancestry, rather than age, CHIP status, or any other factor (Supplementary Figures 1-3). In alignment with what has previously been found in the literature,^59^ ancestry by far influenced methylation variance the most (Figure 1D), and could thus play a major role in altering CHIP development, and its downstream effects, both through epigenetic effects and frequency of CHIP genes mutated.

### Identifying Endotype-Specific Effects on Regulatory Machinery

We first sought to investigate whether CHIP mutations, particularly those in genome stability CHIP driver genes, affected genome mutation burden, both across the whole genome and in CHIP loci. Analysis of whole genome somatic mutation burden showed that CHIP samples (2,380,715.3 mutations/ sample) had higher (but nonsignificant) mutation burden than non-CHIP samples (2,335,504.7, p-value = 0.120) (Figure 2A, Supplementary Table 3). CHIP mutations in genes that participate in genome stabilization (2,381,011.1, p-value = 0.0570), the cohesin complex (2,382,005.1, p-value = 0.0530), and *STAG2* (2,391,950.4, p-value = 0.0740) had marginally significant increases in mutation burden (Figure 2B). We next considered that these CHIP driver genes could be inducing downstream mutations in specific loci that are known to be tolerant to somatic variation, so we estimated total mutation counts within CHIP driver gene loci as known examples (Supplementary Table 4). CHIP samples (6009.8 mutations/ sample) had significantly higher per-sample downstream mutation averages within CHIP genes than non-CHIP samples (5796.5, p-value = 0.0147), with an even higher downstream mutation count in large clones (6059.4, p-value = 0.00670) (Figure 2C). CHIP driver mutations in aggregate within other genome stability CHIP genes did not induce higher downstream mutation counts (5861.2, p-value = 0.259), while only *STAG2* CHIP (6105.0, p-value = 0.0175) had a significantly higher downstream mutation count while other CHIP samples (5954.8, p-value = 0.120) did not (Figure 2D). This suggests the induction of downstream mutations appears to be an effect unique to *STAG2*. Furthermore, *STAG2* was also associated with a higher average count of downstream CHIP driver mutations (1.57 mutations/ sample) that passed the whitelist filter compared to other CHIP samples (1.22 mutations/ sample, p-value = 0.00520), with the effect being greater in large clone *STAG2* samples (1.67 mutations/ sample, p-value = 0.00510) (Figure 2E). *STAG2*, and the genome stabilizing genes in general, do not cause drastic global changes in methylation (mean delta beta: −0.000694, p-value = 0.572) or RNA expression (mean LogFC: 2.831×10^-11, p-value = 3.1×10^-4) compared to other CHIP samples. These findings, along with *STAG2*’s stark effects on downstream mutation burden, indicate that *STAG2* is more likely to act as a “first-hit” in inducing other CHIP mutations that might interfere with cell processes, rather than imparting effects that alter cell function itself.^60^

**Figure 2:**
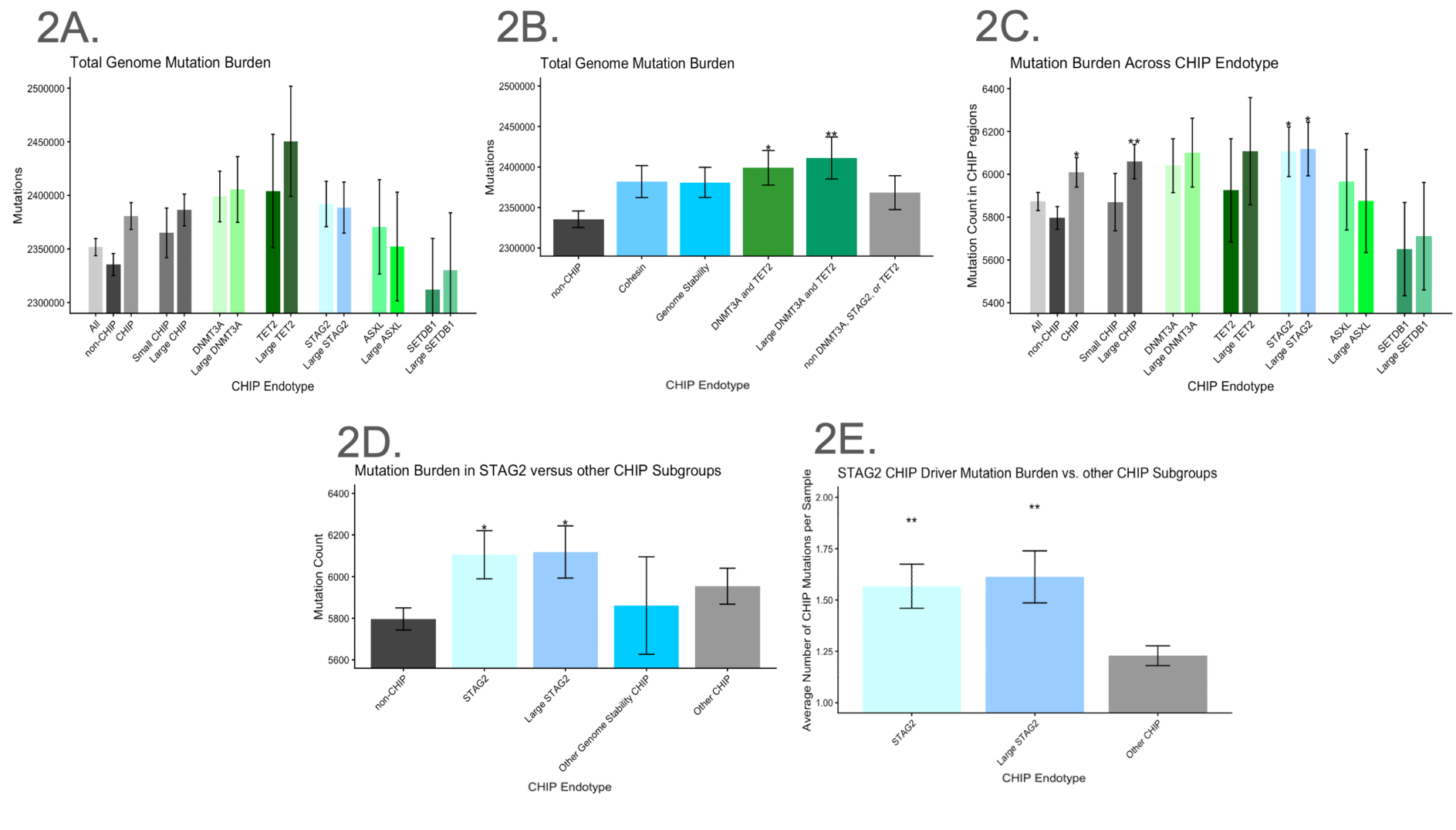
*STAG2* CHIP Drives Genome Instability that Induces further CHIP Mutations **a**, Mutation burden across the whole genome was counted against the Homo_sapiens_assembly38.fasta reference genome from GATK. CHIP driver genes were then grouped by function (Cohesin complex, genome stability, and methylation). For all bar charts, 95% CI error bars are indicated. * indicates a p value < 0.05, and ** indicates a p value < 0.01. **b**, Mutation burden across the genome regions was also counted for the Cohesin complex, genome stability CHIP, and CHIP genes that affect DNA methylation. **c**, Counting of mutation burden across CHIP regions as defined by Vlasschaert et al. (*Blood*, 2023), stratified by CHIP endotype. **d**, Counting of mutation burden across CHIP regions comparing *STAG2* to other CHIP endotypes. **e**, Total CHIP mutations were counted based on mutations in CHIP regions that also passed the whitelist filter provided by Vlasschaert et al. (*Blood*, 2023).

We next sought to determine whether these genetic alterations were accompanied by distinct epigenetic changes through analysis of whole blood DNA methylation. Non-significant global hypomethylation was observed in all CHIP samples compared to participants without CHIP (59.13% hypomethylated, mean delta beta: −0.000869, KS statistic: 0.0406, p-value: 0.191) (Supplementary Figure 4), and even in large clones (55.36% hypomethylated, mean delta beta: −0.00127, KS statistic: 0.0393, p-value: 0.183) (Figures 3A and B, Supplementary Table 5). Much of this global hypomethylation was driven by *DNMT3A* CHIP mutations (60.67% hypomethylated, mean delta beta: −0.00185, KS statistic: 0.0877, p-value: 0.0912), yet when *DNMT3A* mutated samples were removed from the analysis, global hypomethylation was still present (54.73% hypomethylated, mean delta beta: −0.000491, KS statistic: 0.00898, p-value: 0.473) (Figures 3C and D). Similarly, the strength of hypomethylation was greater in large clone *DNMT3A* (65.02% hypomethylated, mean delta beta: −0.00316, KS statistic: 0.0884, p-value: 0.0352) compared to small clones (45.64% hypomethylated, mean delta beta: 0.000609, KS statistic: 0.0354, p-value: 0.125) (Figures 3C and D). Quantile-quantile (QQ) plots demonstrate the larger deviation from the reference probability distribution for DNMT3A CHIP versus all CHIP, and especially *STAG2* CHIP (KS statistic: 0.0237, p-value: 0.706). (Figure 3E). As expected, the global hypomethylation observed across all CHIP samples is accompanied by global increased RNA expression in CHIP overall (mean LogFC: 5.990×10^-11, p-value = 1.0×10^-6) (Figure 3F). However, the strong hypomethylation in *DNMT3A* was not higher than CHIP overall (mean LogFC: 2.071×10^-11, p-value = 0.00840) (Figure 3F). Finally, Horvath clock analysis showed that CHIP (0.616 years, 95%CI: −0.089 − 1.332, p-value = 0.0187), and large clone CHIP (0.710 years, 95%CI: −0.185 - 1.611, p-value = 0.0334), were associated with higher biological age than chronological age (Supplementary Table 6, and Supplementary Figure 5). DNMT3A (1.72 years, 95%CI: 0.318 - 3.117, p-value = 0.00612) and the ASXL family (2.60 years, 95%CI: 0.440 - 4.757, p-value = 0.0132) were the only CHIP endotypes that accelerated biological age significantly. The large clones for *DNMT3A* and the ASXL family CHIP accelerated age even faster, at 2.69 (95%CI: 0.846 - 4.539, p-value = 0.00248) and 2.99 (95%CI: 0.679 - 5.303, p-value = 0.0110) years respectively (Supplementary Table 7).

**Figure 3:**
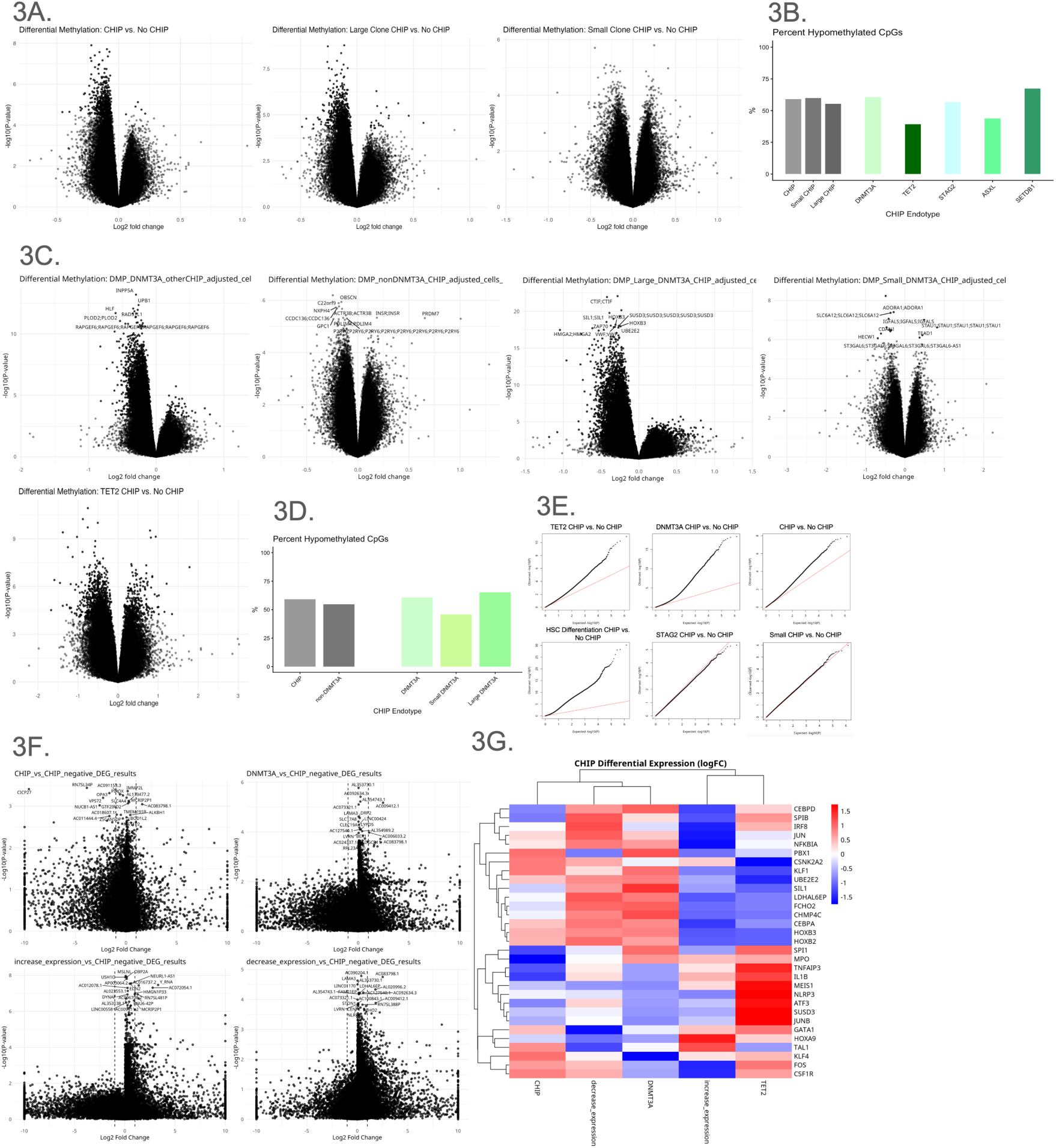
Mutations in CHIP Driver Genes Responsible for Epigenetic Modulation Alter Aging Signatures. **a**, Volcano plots showing differential methylation between CHIP and non-CHIP, stratified by clone size. **b**, Graph showing percent of CpGs that were hypomethylated in different CHIP endotypes. **c**, Volcano plots showing differential methylation between *DNMT3A* CHIP, non *DNMT3A* CHIP, and *TET2* CHIP. **d**, Graph showing percent of CpGs that were hypomethylated in DNMT3A CHIP versus non-DNMT3A CHIP. **e**, Quantile-quantile plots for *TET2* CHIP, *DNMT3A* CHIP, all CHIP, HSC Differentiation CHIP, and *STAG2* CHIP, and small clone CHIP demonstrate deviation form a reference probability distribution. The red line denotes the reference probability distribution. **f**, Volcano plots showing differential RNA expression between CHIP and non-CHIP, *DNMT3A* CHIP, CHIP genes that increase expression, and CHIP genes that decrease expression. **g**, Heatmap showing differential expression of genes between *DNMT3A* and *TET2* CHIP that were significantly hyper or hypomethylated or significantly over or under-expressed in *DNMT3A* and *TET2* CHIP. Genes and CHIP endotypes were grouped by similarity in expression signatures.

Mutations in *TET2* induce more global hypermethylation (60.67% hypermethylated, mean delta beta: 0.00111, KS statistic: 0.0742, p-value: 0.0402), but do not universally impart opposing effects on DNA methylation or expression of genes compared to *DNMT3A* (Figure 3A and B). Despite having a high mean delta beta, mutated *TET2* samples did not appear to have as many significantly hypermethylated CpGs, while still having many significantly hypomethylated CpGs. *TET2* mutations were associated with decreased mean RNA expression (mean LogFC: −1.159×10^-10, p-value = 0.0310). For the most significantly differentially expressed genes in both *DNMT3A* and *TET2*, RNA expression changes were not always opposing (Figure 3G). Furthermore, despite global hypomethylation in *DNMT3A* CHIP and global hypermethylation in *TET2* CHIP, RNA expression could still be decreased in *DNMT3A* CHIP and increased in *TET2* CHIP. This could suggest that other mechanisms besides global methylation are regulating RNA expression in *DNMT3A* and *TET2* samples. Nonetheless, this complicated, not fully discernible relationship between *DNMT3A* and *TET2* reflects others’ findings that *DNMT3A* and *TET2* can act in similar mechanisms in some contexts,^61–63^ and opposite mechanisms in other contexts.^64–68^

### Examining How Systemic Phenotypic Adaptive Responses Change in Certain CHIP Endotypes

Certain CHIP driver genes participate less in multi-omic regulation, but rather in signaling and growth regulation, which we labeled as “phenotypic adaptation.” Having characterized the epigenetic consequences of CHIP-associated mutations, we next investigated their downstream effects on the transcriptome through alternative RNA splicing. RNA splice counts were obtained next to identify if CHIP mutations in RNA spliceosome genes altered splicing function. As a fraction of total RNA reads, CHIP (0.329 splice fraction) did not have a significantly higher splice count than non-CHIP samples (0.319, p-value = 0.112), and large clone CHIP samples (0.331, p-value = 0.0868) did not have significantly higher splice counts (Figure 4A, Supplementary Table 8). When analyzed together, all CHIP driver mutations that affect RNA splicing resulted in a significant increase in RNA splice fraction (0.345, p-value = 0.0364), with an even higher fraction (0.352, p-value = 0.0136) in large clones. However, when further stratified, *SF3A1*, *SRSF2*, and *PRPF40B*, which all regulate RNA splicing activity significantly increased splice fraction (0.370, p-value = 0.0059) while *SF3B1* and *ZRSR2*, which are core proteins in spliceosomes, significantly decreased splice fraction (0.267, p-value = 0.0370) (Figure 4B). Interestingly, the effects of ASXL (0.338, p-value = 0.00680) and large ASXL (0.339, p-value = 0.0189) samples on RNA splice fraction were significant, while only one participant had concurrent mutations in an ASXL and RNA splicing-associated CHIP gene.

**Figure 4:**
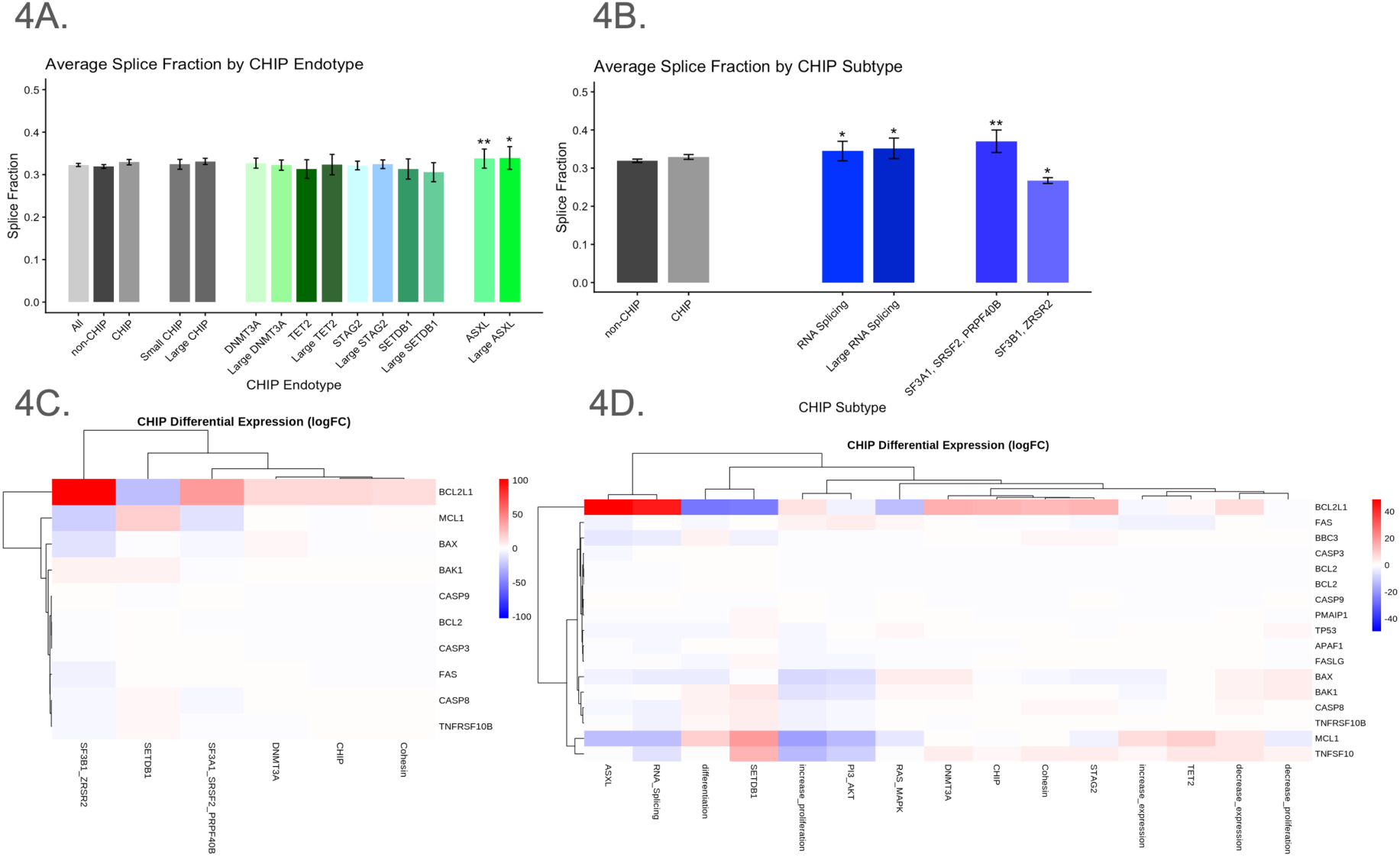
Mutations in CHIP Genes that Regulate RNA Splicing Reduce Apoptotic Response. **a**, Average splice fraction was calculated from RNA expression analyses by dividing number of spliced reads by total number of RNA reads. A significantly higher or lower splice fraction could thus indicate splicing dysfunction. For all bar charts, 95% CI error bars are indicated. * indicates a p value < 0.05, and ** indicates a p value < 0.01. **b**, Splice fractions were then calculated for CHIP driver genes that participate in RNA splicing, and their large clones. RNA splicing CHIP was then further stratified into genes that make up the spliceosome structure (*SF3B1*, *ZRSR2*) and genes that are involved in splicing regulation (*SF3A1*, *SRSF2*, *PRPF40B*). **c**, Heatmap shows the similar over-expression of anti-apoptotic factors and downregulation of apoptotic factors in both genes involved in spliceosome structure and splicing regulation. Scale is **d**, Heatmap shows similar regulation of apoptotic factors in ASXL family CHIP that is not present in other CHIP endotypes. Genes and CHIP endotypes were grouped by similarity in expression signatures.

Given the strong association between RNA splicing and cell growth and death,^69^ we sought to investigate if apoptosis was affected in RNA splicing CHIP samples. Despite having differing effects on RNA splice fractions, *SF3A1, SRSF2, PRPF40B, SF3B1, ZRSR2*, and the ASXL genes displayed strong overexpression of the *BCL2L1* gene (mean LogFC: 44.032, p-value = 0.000310), a major apoptotic factor that has two alternatively spliced forms-a long form that is anti-apoptotic, and a short form that is pro apoptosis (Figures 4C and D). We propose that CHIP mutations in RNA splicing genes, although they affect splicing differently, create enough dysregulation in RNA splicing machinery that apoptosis is inhibited, and mutant cells are allowed to proliferate. The upregulation of *BCL2L1* and anti-apoptotic machinery is surprising in the ASXL CHIP mutations, as ASXL mutations are typically pro-apoptotic.^70,71^

Another category of CHIP driver genes that affect phenotypic adaptation were genes that participate in HSC differentiation of different hematologic cell types. CHIP driver genes in this category include *CSF3R* (macrophages and granulocytes), *IKZF1* (lymphocytes (B- and T-cell)), *IKZF2* (regulatory T cells), and *RUNX1* (hematopoiesis). We hypothesized that dysregulation of differentiation would alter cell-type proportions and have more specific functional effects than the previously mentioned multiomic regulatory CHIP driver genes.

We found that HSC differentiation CHIP mutations resulted in the greatest hypomethylation (58.19% hypomethylated, mean delta beta: −0.00468, KS statistic: 0.240, p-value: 0.0498) of all CHIP endotypes. Even so, HSC differentiation CHIP mutations were also associated with significantly hypermethylated CpGs as well (7154 significantly hypermethylated, 5900 significantly hypomethylated) (Figure 5A). This is unsurprising, as differentiation of stem cells is driven largely by epigenetic signals.^72^ From our methylation data, cell–type proportions could be estimated, and between CHIP endotypes remained relatively consistent (Figure 5C). However, mutations that alter HSC differentiation resulted in a higher estimated percentage of CD8+ T cells (11.06%, p-value = 0.0508), compared to other non-CHIP (6.84%) and other CHIP (6.85%, p-value = 0.998) samples (Figure 5B, Supplementary Table 9). Unlike Horvath age and mutation burden, it does not appear that other endotypes of CHIP affect CD8+ T cell count at all. Large clone HSC differentiation CHIP driver mutations were associated with even higher CD8+ T cell proliferation (14.47%, p-value = 0.0139). Interestingly, it does not appear any specific cell type was concordantly reduced as a result of the higher CD8+ T cell count, and no other forms of CHIP appeared to alter cell type proportions in any significant way (Figure 5C).

**Figure 5:**
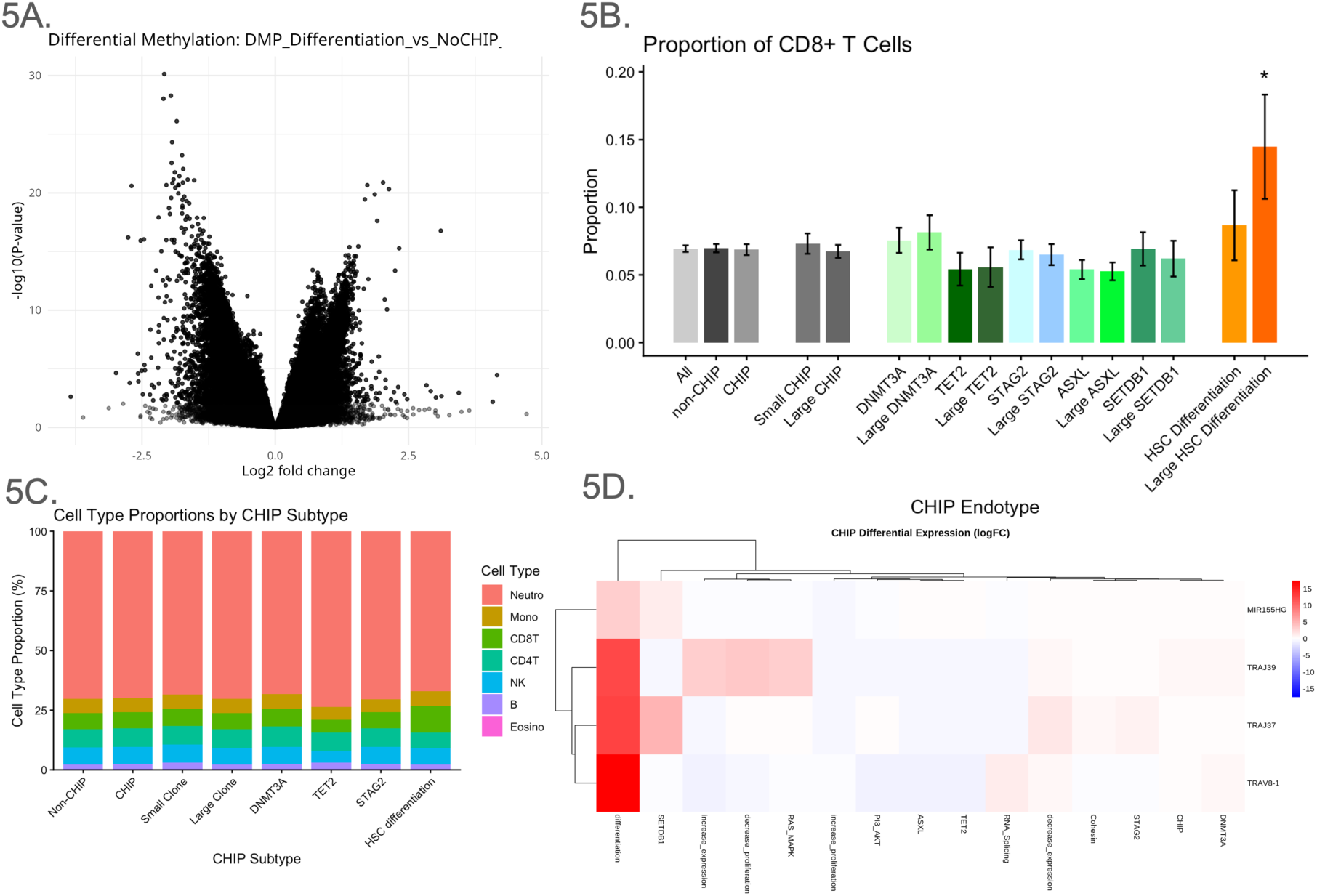
CHIP Mutations that Alter Hematopoietic Stem Cell Differentiation Increase Inflammation through Adaptive Immune Response Pathways. **a**, Volcano plot showing differential methylation between HSC differentiation CHIP, and non-CHIP. **b**, Cell type proportions (for CD4⁺ T cells, CD8⁺ T cells, B cells, NK cells, monocytes, neutrophils, and eosinophils) were estimated with the Houseman deconvolution method through the EpiDISH R package. CD8+ T cells in HSC differentiation CHIP were the only cell-type that was differentially represented in any CHIP endotype. 95% CI error bars are indicated. * indicates a p value < 0.05, and ** indicates a p value < 0.01. **c**, Cell type proportions for CD4⁺ T cells, CD8⁺ T cells, B cells, NK cells, monocytes, neutrophils, and eosinophils for common CHIP endotypes. **d**, Heatmap showing differential expression of key genes involved in adaptive immune response in HSC differentiation CHIP versus other CHIP endotypes. Genes and CHIP endotypes were grouped by similarity in expression signatures. CH-all clonal hematopoiesis samples, inc_exp-CH genes that epigenetically increase RNA expression, dec_exp-CH genes that epigenetically decrease RNA expression, ASXL-ASXL family genes, RAS-genes involved in the RAS/MAPK pathway, PI3-genes involved in the PI3K/AKT pathway, inc_prolif-CH genes that increase cell proliferation, dec_prolif-CH genes that decrease cell proliferation, Diff-CH genes involved in HSC differentiation, RNA-CH genes involved in RNA splicing.

Additionally, GO Enrichment analysis of RNA expression revealed that the only significantly enriched biological pathway was “adaptive immune response”. Some of the genes with increased expression in the adaptive immune response pathway included *TNFRSF9* (mean LogFC: 1.7845, p-value = 5.696×10^-4), *TRAV8-1* (mean LogFC: 17.440, p-value = 2.055×10^-12), *TRAJ39* (mean LogFC: 12.481, p-value = 3.566×10^-7), *TRAJ37* (mean LogFC: 12.843, p-value = 1.932×10^-5), and *MIR155HG* (mean LogFC: 3.151, p-value = 0.0224), and these genes were visibly less upregulated in other CHIP endotypes (Figure 5D). *TNFRSF9* is highly expressed on T-cells. *TRAV8-1, TRAJ39*, and *TRAJ37* are all expressed on T-cell receptors, and *MIR155HG* mounts optimal primary and memory responses by suppressing inhibitory signaling proteins, which enhances the T cell proliferation.

We then asked whether this proliferation of CD8+ T cells and enriched adaptive immune response altered inflammation in any meaningful way. *CCL5* (mean LogFC: 77.148, p-value = 0.0197) and *S100A9* (mean LogFC: 2744.638, p-value = 0.0420) were strongly and uniquely upregulated in HSC Differentiation CHIP samples (Figure 5D). *JUNB* is a master regulator of cell proliferation, tissue homeostasis, and immune system responses, and *CCL5* is a pro-inflammatory cytokine. *CCL5* is highly expressed by activated CD8+ T cells, cytotoxic T cells, and natural killer cells. *S100A9* is a pro-inflammatory protein primarily produced by neutrophils and monocytes that is involved in regulating immune responses, cell migration, and antimicrobial activities.^73^ *S100A9* also primarily forms the inflammatory calprotectin heterodimer with *S100A8*, which was also upregulated (mean LogFC: 223.972, p-value = 0.0340).

These findings suggest that HSC differentiation CHIP mutations do not just uniformly induce hematopoiesis, but might drive CD8+ T cell proliferation, which triggers the adaptive immune response and induces general inflammation that has been seen canonically in CHIP patients. This differs from much of the literature which reports that CHIP induces innate inflammatory activation,^15,74,75^ whereas the genes in HSC Differentiation CHIP seem more likely to drive adaptive and cytotoxic immune response.

The final major CHIP endotype was proliferative signalling, which we also labeled as a “phenotypic adaptation.” Similar to HSC differentiation, we suspected proliferative signalling CHIP mutations would have specific rather than broad functional effects, particularly on biological pathways downstream of the CHIP driver gene. Thus, we focus much of our discussion on these related pathways as well as specific genes that were significantly up or downregulated in our analyses. Analysis of RNA expression of CHIP mutations in driver genes that increase cell proliferation showed GO enrichment of response to wounding (p-value = 0.0367) and wound healing (p-value = 0.0357) (Figure 6A). Specific genes in these pathways that were differentially expressed include: *PF4V1* (mean LogFC: 22.5366, p-value = 2.397×10^-7)*, PF4* (mean LogFC: 117.766, p-value = 2.154×10^-4)*, GP1BB* (mean LogFC: 237.632, p-value = 2.827×10^-5)*, ITGA2B* (mean LogFC: 44.978, p-value = 1.916×10^-6)*, TREML1* (mean LogFC: 40.834, p-value = 4.183×10^-6)*, MYL9* (mean LogFC: 34.797, p-value = 5.651×10^-6) (Figure 6B). Upregulation of these genes was unique to increased cell proliferation CHIP, and these genes were largely downregulated in CHIP genes that decrease proliferative signalling (Figure 6C).

**Figure 6:**
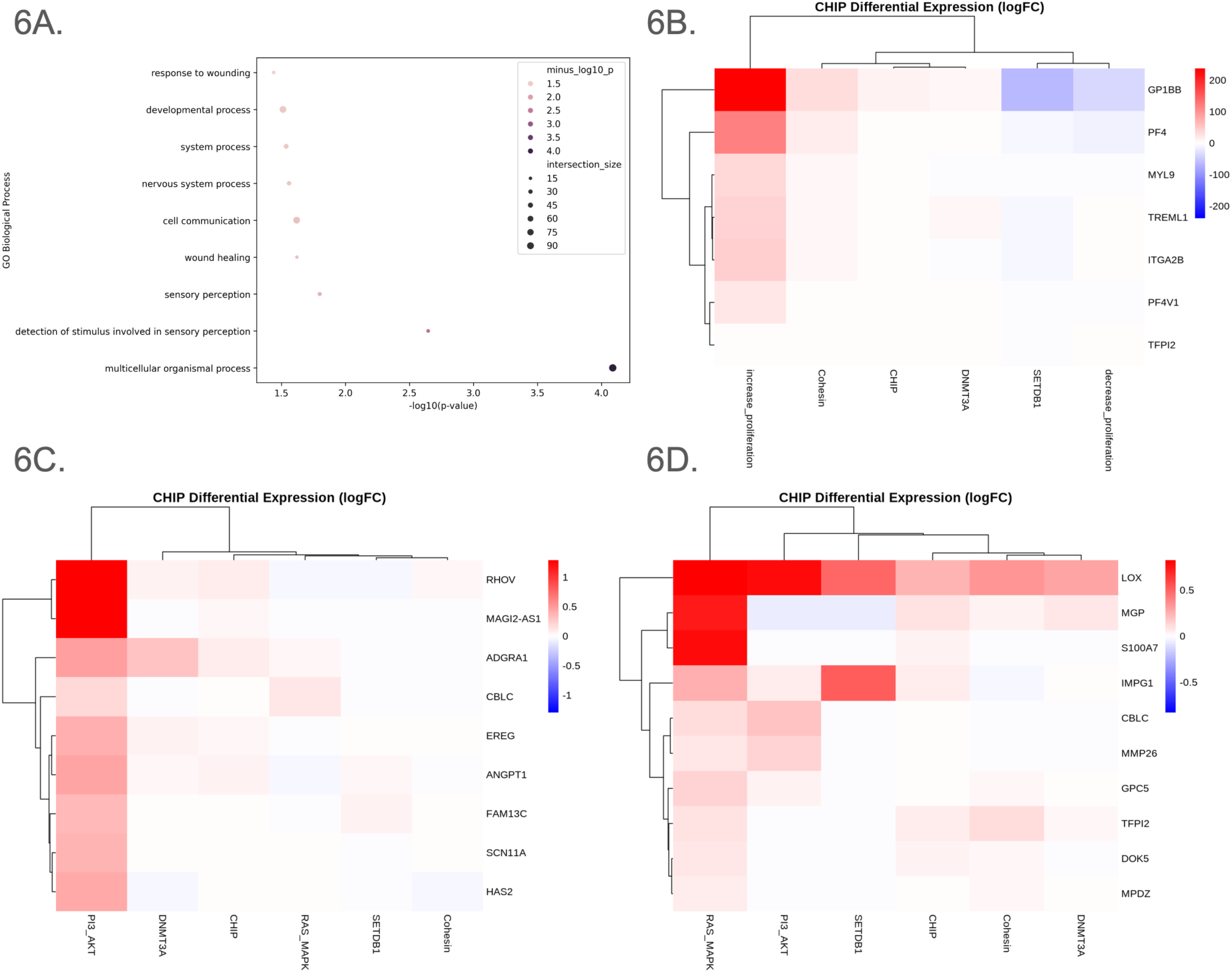
Mutations in CHIP Driver Genes Involved in Metabolic or Proliferative Signaling Specifically Affect These Pathways. **a**, GO enrichment analysis shows hemostasis, coagulation, wound healing, and cell adhesion were all significantly enriched biological pathways in CHIP driver genes that increase proliferation compared to non-CHIP samples. **b**, Heatmap shows upregulation of genes involved in the hemostasis, coagulation, and wound healing in CHIP driver genes that increase proliferation but not other CHIP endotypes. **c**, Heatmap shows genes significantly upregulated in the PI3K/AKT pathway are not upregulated in other CHIP endotypes. **d**, Heatmap shows genes significantly upregulated in the Ras/MAPK pathway are not upregulated in other CHIP endotypes.

Both the PI3K/AKT and Ras/MAPK CHIP endotypes had unique genes that participate in their respective pathways that were significantly upregulated but not differentially expressed in other CHIP samples. *RHOV* (mean LogFC: 1.283, p-value = 2.627×10^-7) and *MAGI2-AS1* (mean LogFC: 1.290, p-value = 1.205×10^-8) are strongly upregulated in PI3K/AKT but not in other CHIP samples or in Ras/MAPK (Figures 6D and E). The altered expression of both genes have been implicated in tumor pathogenesis.^76,77^ *LOX* (mean LogFC: 0.825, p-value = 0.0360) and *MGP* (mean LogFC: 0.733, p-value = 0.0133) are strongly upregulated in Ras/MAPK samples, but less so in other CHIP or PI3K/AKT (Figures 6F and G), both of which also play roles in oncological development.^78,79^ Despite all affecting tumorigenesis, these examples most clearly demonstrate CHIP mutations having consistent opposite effects on downstream pathways, and support the suggestion that not only is stratification by CHIP driver gene beneficial to make any claims about downstream effects of CHIP, but also necessary due to the opposing functional nature of many CHIP driver genes, and the resulting divergent effects of CHIP mutations.

## Discussion

Prior literature has shown that inflammation is one of the primary phenotypic hallmarks of CHIP.^10,15,16,18,19^ Our stratifications of CHIP indicate that this inflammation is not uniform between CHIP endotypes. Strongly divergent effects on RNA expression were observed in innate immune response (Figure 7A), adaptive immune response (Figures 7B-E), chemokine (Figure 7F) and interferon expression (Figure 7G), monocyte activation (Figure 7H), and allergic response (Figure 7I). For example, the *PYCARD* gene, modulator of innate immune responses and pyroptosis, was strongly upregulated in some endotypes and strongly downregulated on other endotypes. This downregulation of innate immunity counters what has previously been observed in CHIP.^74,75^ Similar bidirectional effects were observed in each of the other immune programs. Each CHIP endotype appeared to have its own unique signature, as if one gene was unidirectional between two CHIP endotypes, it does not mean that other genes were also unidirectional. For example, *DNMT3A* and *TET2* typically had opposing effects on each immune response, except for eosinophils. *DNMT3A* increased Th1 signaling and decreased Th2 signaling, while *TET2* increased Th2 signaling and decreased Th1 signaling. Other CHIP endotypes, such as HSC Differentiation, and PI3K/AKT and Ras/MAPK, unidirectionally affect Th1 and Th2 signaling, possibly bypassing the counteractive regulatory mechanisms between Th1 and Th2. Even within particular immune programs, such as the innate immune and TReg responses, RNA expression directionality was not uniform, suggesting a more complex relationship between certain CHIP endotypes and these immune programs beyond just up or downregulation.

**Figure 7:**
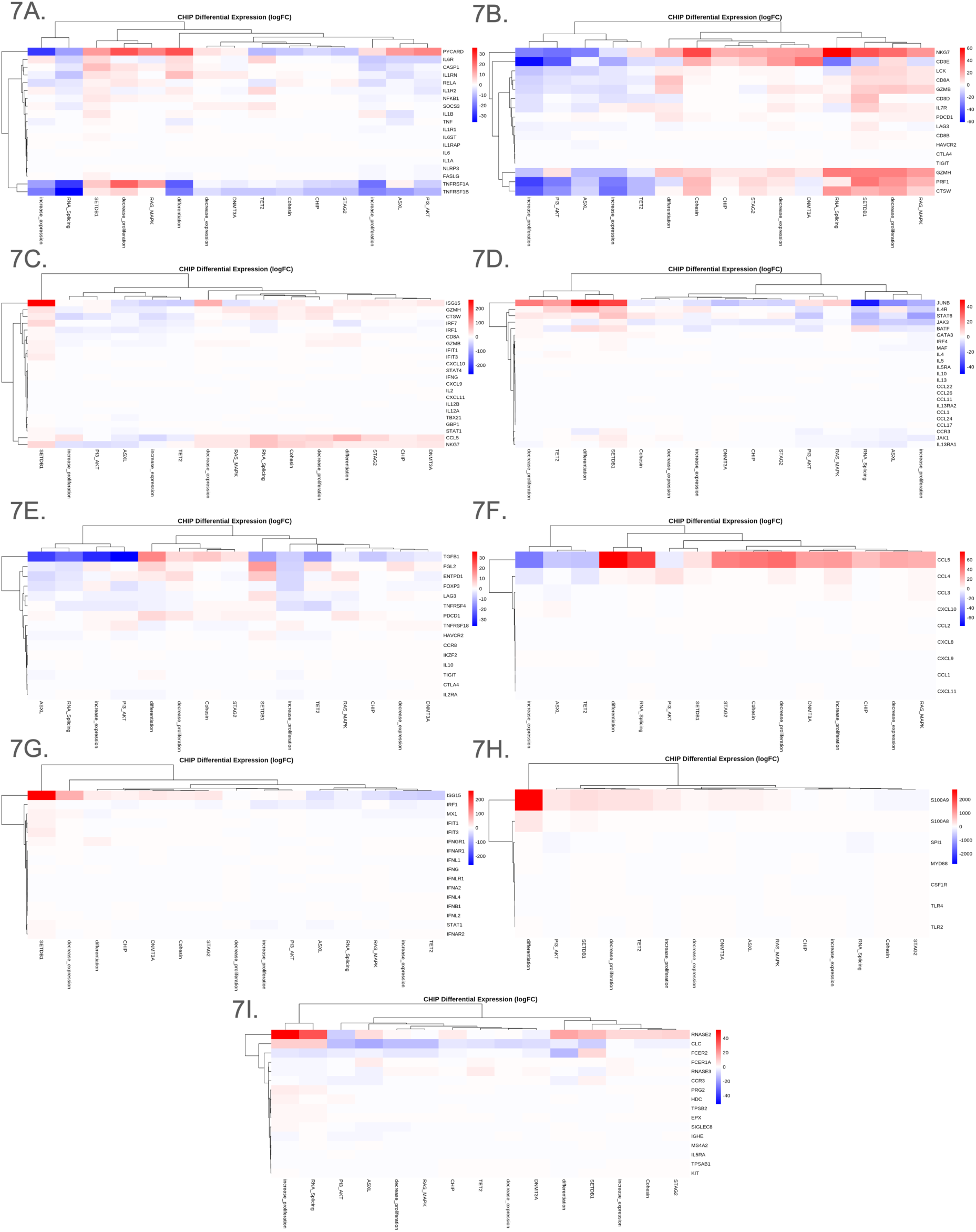
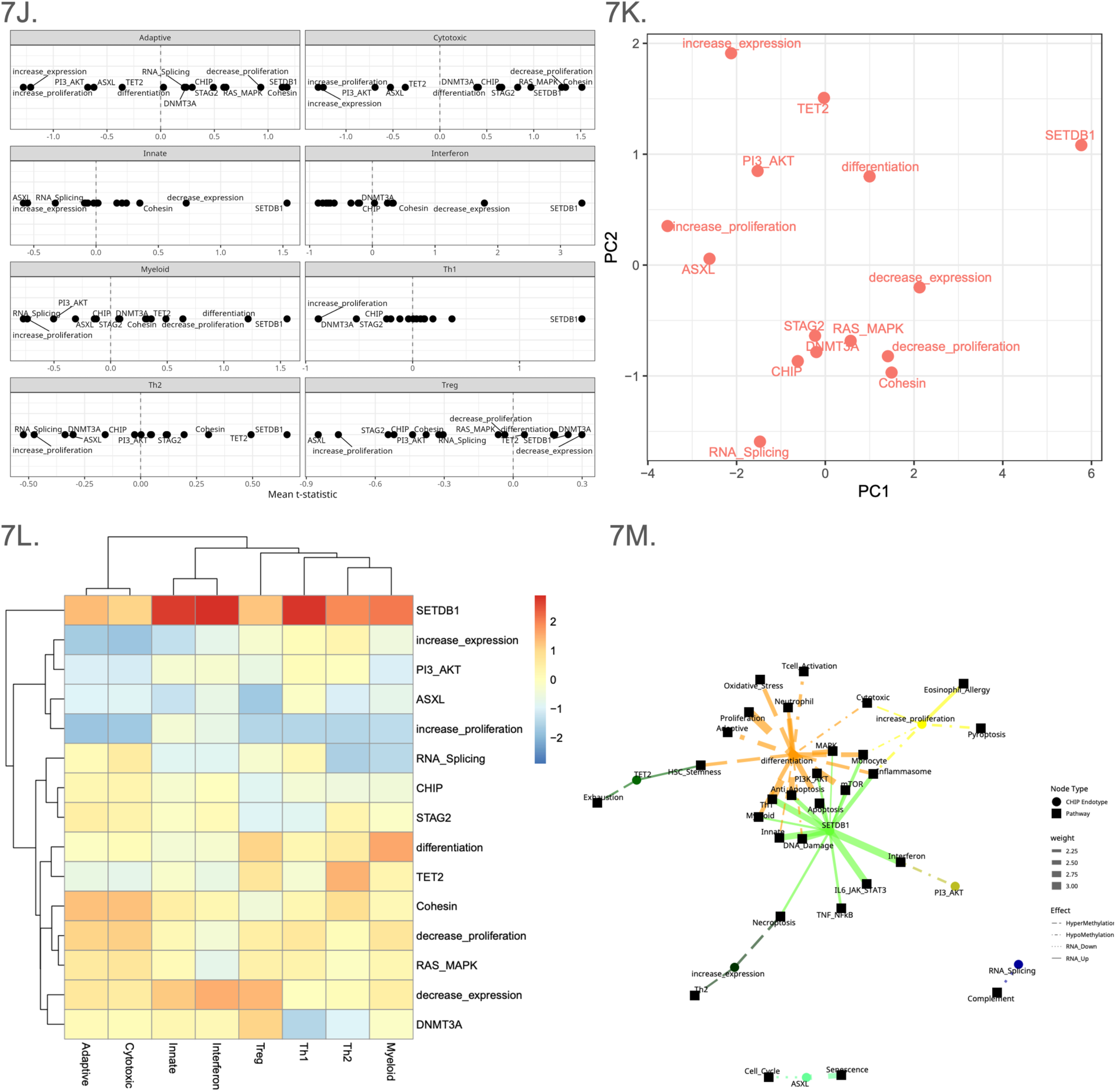
CHIP as a Whole Lacks a Driving Mechanism, and its Endotypes Induce Unique Downstream Inflammatory Effects. **a-i**, Heatmaps show consistent effects on various inflammatory pathways (innate immune response, T cell activation, Th1 signaling, Th2 signaling, TReg signaling, chemokines, interferons, monocyte activation, and eosinophils/ allergic response) depending on CHIP endotypes. CHIP-all clonal hematopoiesis of indeterminate potential samples, inc_exp-CHIP genes that epigenetically increase RNA expression, dec_exp-CHIP genes that epigenetically decrease RNA expression, ASXL-ASXL family genes, RAS-genes involved in the RAS/MAPK pathway, PI3-genes involved in the PI3K/AKT pathway, inc_prolif-CHIP genes that increase cell proliferation, dec_prolif-CHIP genes that decrease cell proliferation, Diff-CHIP genes involved in HSC differentiation, RNA-CHIP genes involved in RNA splicing. **j**, Mean t-statistics were calculate for each isoform and each inflammatory pathway relative to non-CHIP samples. Genes involved in the t-statistics calculations were the same genes used in each heatmap. The dotted vertical line indicates no absolute change in expression of a given pathway. **k**, Each CHIP endotype was clustered based on overall effects of all inflammatory pathways. **l**, Heatmap shows bidirectional regulation of inflammatory pathways based on CHIP endotype. Expression levels of each pathway were scaled as a z-score. CHIP endotypes and immune pathways were grouped by similarity in expression signatures. **m**, Graph showing CHIP endotypes with significant epigenetic or RNA expression associations with immune and cell survival or proliferation pathways.

**Figure 8:**
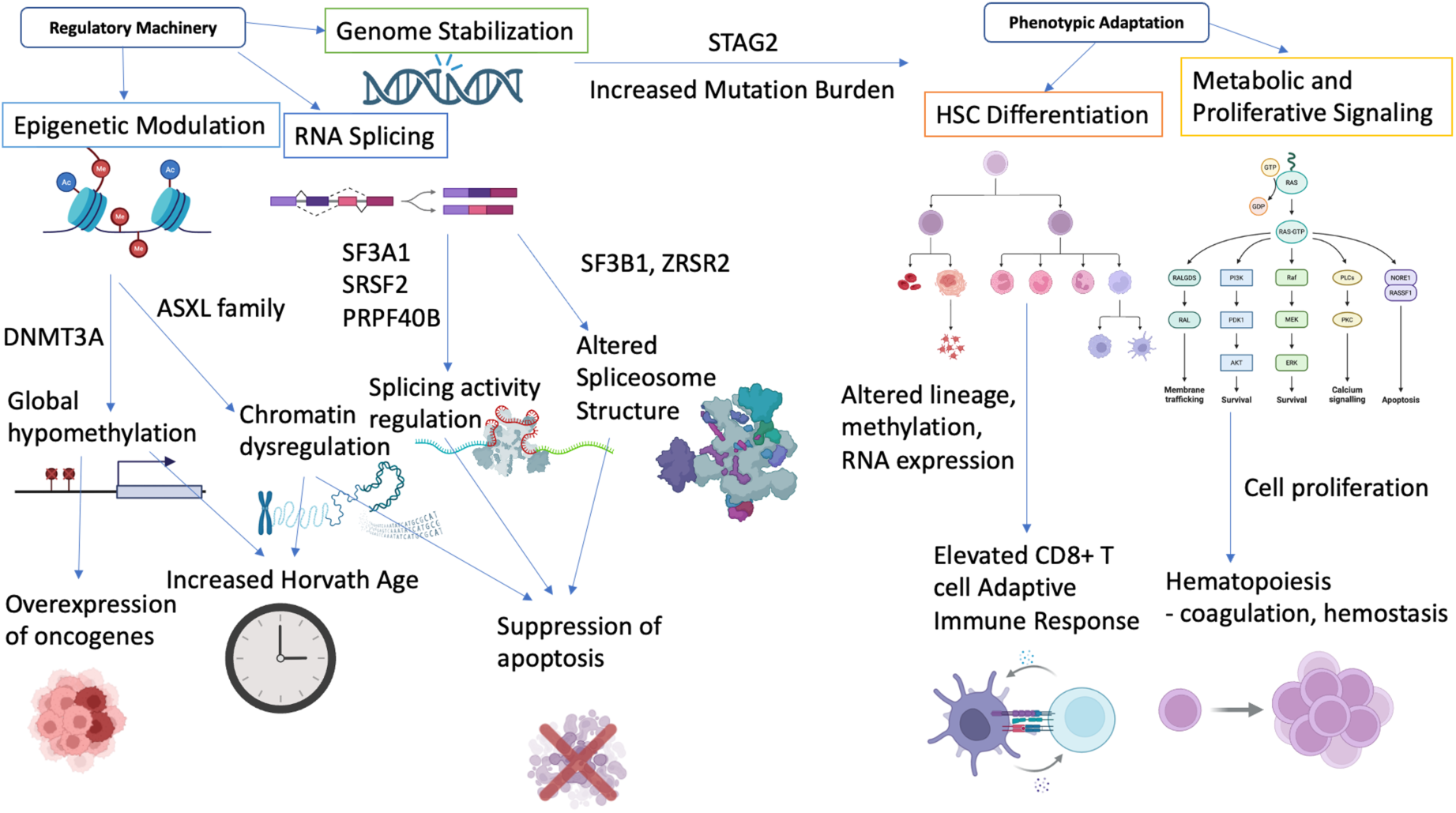
A Proposed Mechanism for CHIP’s Shared and Divergent Effects. Schematic showing a proposed mechanism of CHIP progression depending on CHIP driver gene, indicating functional categorizations, mechanisms of dysregulation, interactions with other CHIP genes, and downstream effects.

Figure 7J shows the overall effects each of the CHIP endotypes have on RNA expression of genes involved in adaptive, cytotoxic, innate, interferon, myeloid, Th1, Th2, and TReg immunity. Interestingly, in most immune programs, *SETDB1* had the highest expression, suggesting a stronger relationship with inflammation than other endotypes. CHIP endotypes with opposing functions, such as *DNMT3A* and *TET2*, increased expression and decreased expression, and increased proliferation and decreased proliferation, tended to cluster away from each other (Figure 7K). Each immune program was upregulated by certain endotypes, and downregulated or unaffected by others (Figure 7L). Combining methylation effects with RNA expression effects, we constructed a graph displaying the relationship CHIP endotypes have with immune programs as well as other major cell survival and proliferation functions (Figure 7M). Notably, mutations in *SETDB1* and HSC differentiation CHIP were linked to more biological pathways. CHIP, *STAG2*, and the Ras/MAPK pathway were not significantly associated with any biological pathways. It was interesting that the innate immune response was upregulated in mutations in CHIP genes that decrease expression, a category that includes *DNMT3A*. CHIP has always been associated with an elevated innate immune response,^15,74,75^ and *DNMT3A* being the most commonly mutated CHIP gene in most other datasets could have biased those results.

In order to better clarify the seemingly paradoxical roles CHIP can play in the progression of diseases such as solid tumors, neurodegenerative disease, and CKD, we stratified CHIP by driver gene function, clone size and ancestry. We found that the distribution of CHIP mutations in non-White samples differed from previous studies where most samples were non-Hispanic white, with *STAG2* being the most commonly mutated CHIP gene in the MAGENTA dataset. In addition, across all effects, it appears the magnitude of CHIP’s effects are dose-dependent, with large VAF CHIP samples exacerbating these effects. This finding is significant because as more research is done on the potential implications of CHIP for overall patient health, the clone size of a CHIP mutation could be a major determinant for disease progression and therefore clinical relevance.

Our results showed that effects of CHIP were not uniform, and these effects were driven by specific driver genes with relevant functions. *DNMT3A* CHIP caused global hypomethylation, while *TET2* CHIP did not conversely drive global hypermethylation. *STAG2* CHIP increased mutation burden particularly in CHIP regions, acting as a “first-hit” for more putative CHIP mutations. RNA splicing CHIP resulted in propagation of anti-apoptotic machinery, potentially contributing to mutant cell proliferation. HSC differentiation CHIP strongly triggered the adaptive immune response and resulted in CD8+ T-cell expansion. CHIP mutations that increased proliferative signaling were uniquely enriched for coagulation and cell adhesion, another canonical hallmark of CHIP. In many of our analyses, such as with global methylation, Horvath age clock, and mutation burden, we found that CHIP appeared to have significantly different results compared to non-CHIP participants. However, when stratified by certain subtypes of CHIP, there were often only a few or one CHIP subtype(s) that were significantly different, and the rest of CHIP more closely resembled the non-CHIP samples. This not only supports our claim that CHIP stratifications should be used when investigating any relationship CHIP may have with a disease or phenotype, but also that previous association studies could have conflated associations when in reality only a few subtypes of CHIP actually mediated that relationship, but the effects of those subtypes were strong enough for there to seem to be a strong association across all CHIP samples. Importantly, these CHIP isoforms not only induced differences in epigenetic, genome, splicing, and specific pathway machinery that coincide with their canonical function, but also had clear cut divergent effects, particularly in how they induce inflammation. For example, CHIP genes involved in HSC differentiation activated adaptive immune response which significantly differs from how most literature has demonstrated that CHIP’s primary involvement in the immune system is in the innate immune response. CHIP has canonically been shown to increase Th1 signaling, and we show here that CHIP on average leads to a decrease in Th2 signaling. Yet, when stratified further, CHIP isoforms could increase or decrease signaling in either Th1 or Th2 independently of their effects on the other. Finally, when performing RNA expression and GO enrichment analysis on CHIP overall, a lack of significant results implies a lack of any shared mechanisms that CHIP influences, and thus studying CHIP as a single entity becomes incredibly moot.

The results of this paper leave room for additional exploration into both the best ways to stratify CHIP driver mutations and to investigate CHIP’s impact on aging, and other diseases and phenotypes. Although we used a previously established method of calling CHIP, CHIP driver genes continue to be discovered and although unlikely, it is possible that some participants with larger CHIP VAFs could have full blown hematological cancer, which was not accounted for in our analyses and could have its own completely different set of implications. Therefore, we do not claim that our CHIP and no CHIP categorizations are by any means perfectly accurate. Another shortcoming with our CHIP diagnosis pipeline is that it is impossible to know the onset of a CHIP event, and therefore the duration at which participants had CHIP. If participants with a certain CHIP endotype had CHIP for longer, this would likely influence the severity of many of our analyzed outcomes that proliferate over time, particularly the accumulation of somatic mutations.

A larger dataset would also prove useful to better validate the significance of the effects found in this paper. Stratifying CHIP by driver gene function and by families, complexes, or pathways meant that certain endotypes had few samples. Increasing the power of the study could better validate the effects of these pathways. Additionally, not all canonical CHIP driver genes had mutations present in our dataset and were thus not included in the study. Finally, increasing power could also allow for exploration into differing functions of CHIP mutations within genes. Although CHIP mutations are typically loss-of-function, some are gain-of-function,^80,81^ and we did not differentiate between these possible differences in our analyses.

Another shortcoming of the data that we utilized is that it was initially curated for AD research, meaning that a disproportionate percent (51.5%) of participants had an AD diagnosis. Although all our analyses controlled for AD status, the presence of such a high percent of AD participants could have skewed the distribution of CHIP samples or expression data without our knowing. Additionally, exploration into how ancestry impacts distribution of CHIP genes and downstream effects could benefit from a dataset that includes participants from beyond just African, Hispanic, and non-Hispanic White lineages. Finally, the implications of our CHIP stratifications should be investigated particularly in diseases where there is conflicting evidence behind CHIP’s role in those diseases, as this could clarify discrepancies and provide more specific targets for therapeutic discoveries.

Our multi-omics analyses provide a function-based framework for categorizing CHIP through integration of driver gene function, clone size, and ancestry. We observe that CHIP mutations, depending on the driver gene, can impart unique and even opposing effects on DNA methylation, RNA expression, RNA splicing, genome mutation burden, hematological cell type proportions, and inflammatory pathways. We strongly urge further CHIP studies to clearly and deliberately stratify CHIP by driver gene function, and to demonstrate that any global effects of CHIP are truly progressing by the same mechanism, rather than the effect of a single or few CHIP subtypes influencing the results. These categorizations will improve mechanistic interpretation of conflicting or heterogeneous findings surrounding CHIP’s effects on age-related diseases, such as Alzheimer’s disease, cardiovascular disease, and solid tumors.

## Acknowledgements

This work was supported by grants from the National Institutes of Health T32GM152319 (Liu), National Institute on Aging T32AG071474 (Bush), and R01AG070935 (Griswold, Bush).

## Funder Information Declared

National Institutes of Health, R01AG070935

## Notes

### Competing Interest Statement

The authors have declared no competing interest.

