## Supplemental Figures for "Heterogeneous Effects Across Clonal Hematopoiesis of Indeterminate Potential (CHIP) Driver Mutations Challenges a Unified Model of CHIP"

### Supplementary Figure 1: Samples Cluster by Ancestry and Batch

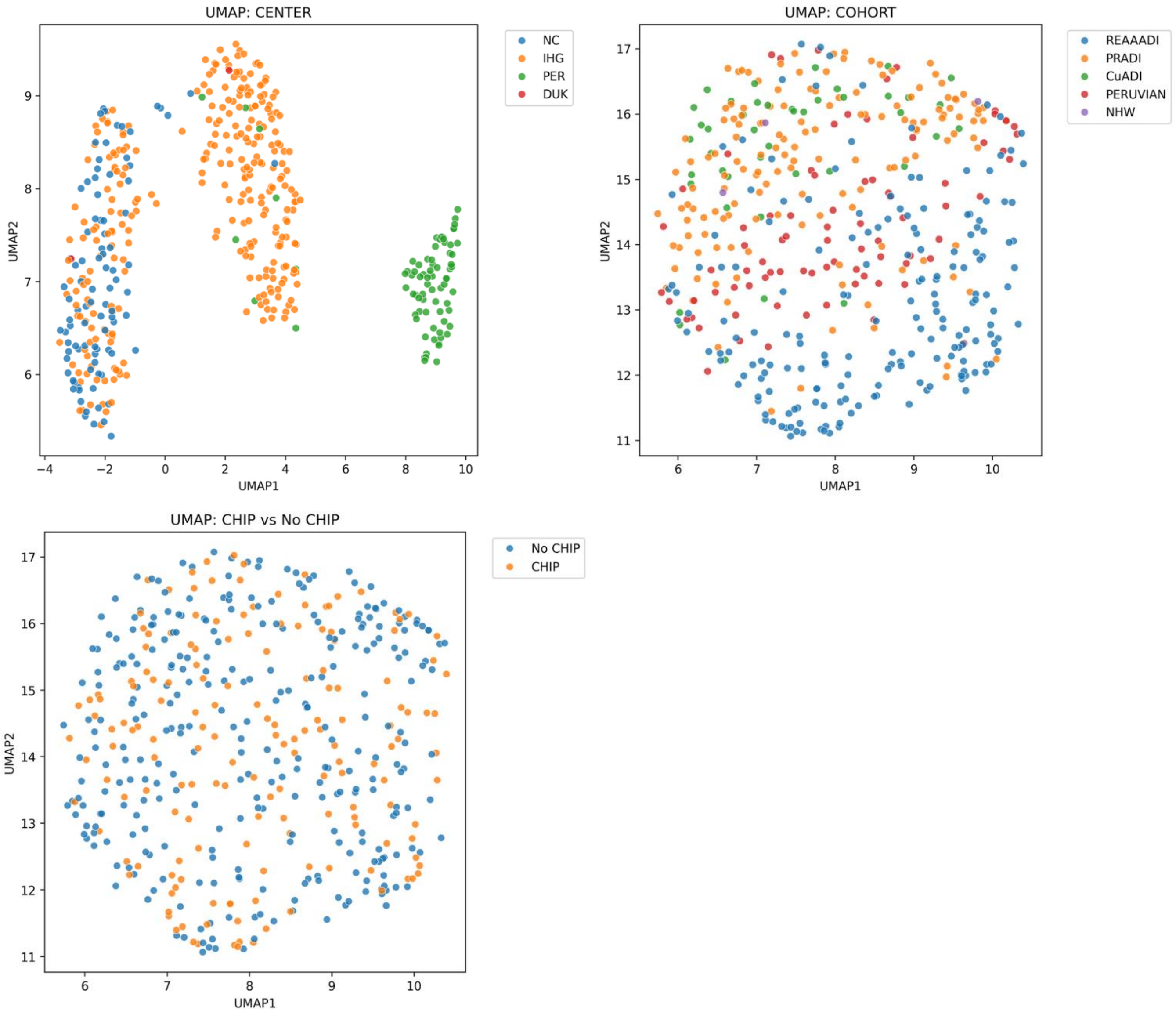

### Supplementary Figure 3: Volcano Plots Differences by Ancestry Will Require Further

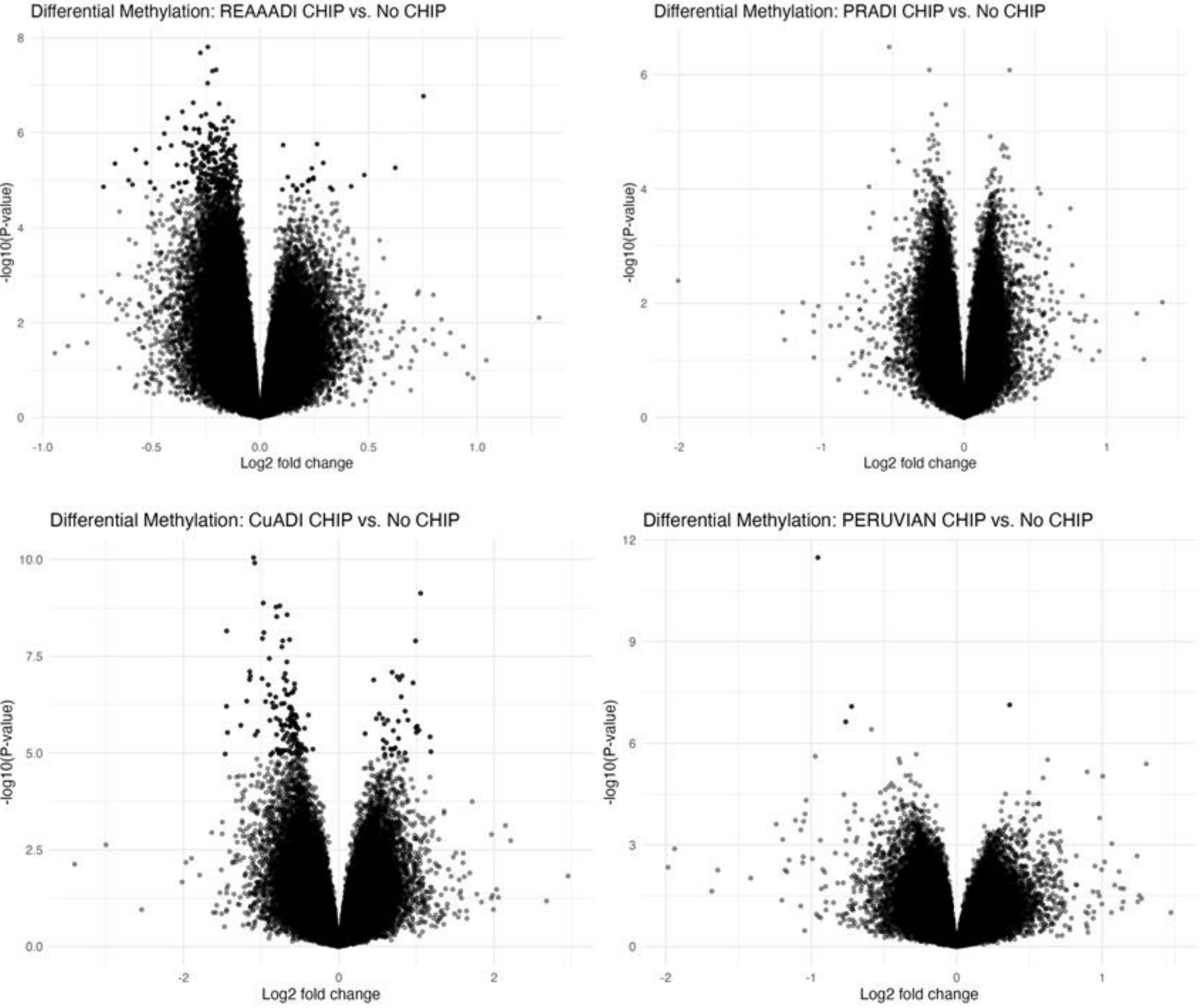

### Supplementary Figure 4

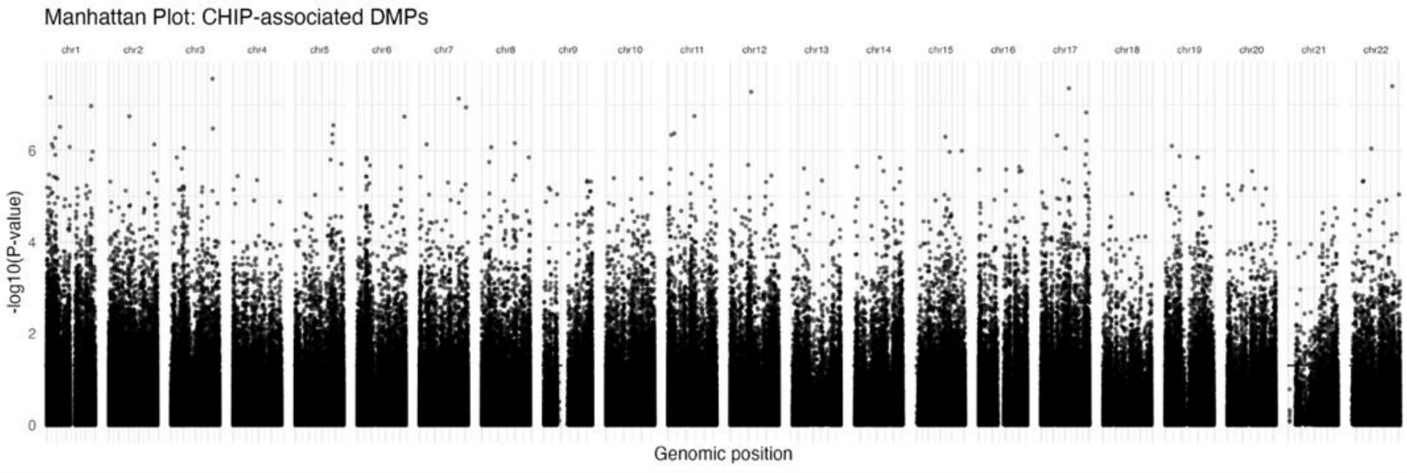

### Supplementary Figure 5

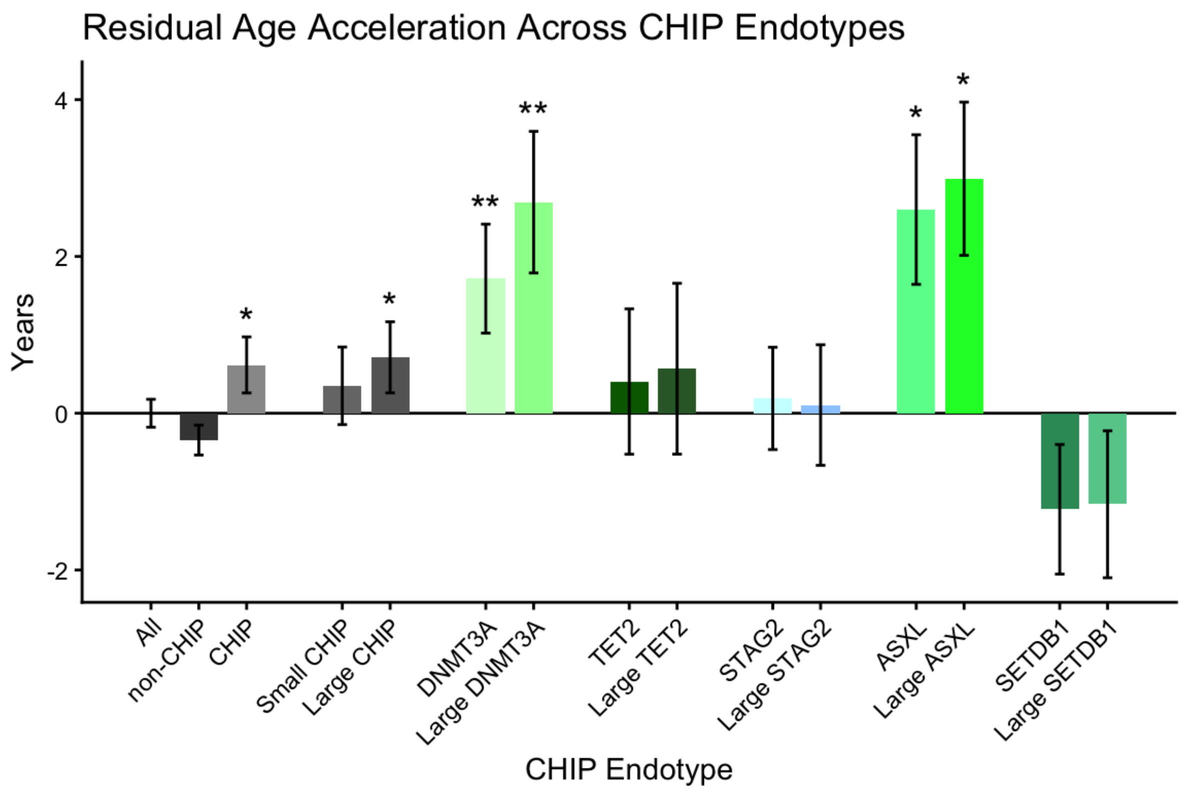


Residual age acceleration was calculated using 353 CpG sites from the Horvath methylation age clock. 95% CI error bars are indicated. * indicates a p value < 0.05, and ** indicates a p value < 0.01.
